# pheno-seq – linking morphological features to gene expression in 3D cell culture systems

**DOI:** 10.1101/311472

**Authors:** Stephan M. Tirier, Jeongbin Park, Friedrich Preußer, Lisa Amrhein, Zuguang Gu, Simon Steiger, Jan-Philipp Mallm, Marcel Waschow, Björn Eismann, Marta Gut, Ivo G. Gut, Karsten Rippe, Matthias Schlesner, Fabian Theis, Christiane Fuchs, Claudia R. Ball, Hanno Glimm, Roland Eils, Christian Conrad

**Affiliations:** Division of Theoretical Bioinformatics, German Cancer Research Center (DKFZ), Heidelberg, Germany; Center for Quantitative Analysis of Molecular and Cellular Biosystems (BioQuant), University of Heidelberg, Heidelberg, Germany; Digital Health Center, Berlin Institute of Health (BIH)/Charité-Universitätsmedizin Berlin, Berlin, Germany; Max Delbrück Center for Molecular Medicine, Berlin Institute for Medical Systems Biology, Berlin, Germany; Helmholtz Zentrum München - German Research Center for Environmental Health, Institute of Computational Biology, Munich, Neuherberg, Germany; Department of Mathematics, Technische Universität München, Munich, Germany; Division of Chromatin Networks, German Cancer Research Center (DKFZ), Heidelberg, Germany; Heidelberg Center for Personalized Oncology, DKFZ-HIPO, DKFZ, Heidelberg, Germany; CNAG-CRG, Centre for Genomic Regulation, Barcelona Institute of Science and Technology, Barcelona, Spain; Universitat Pompeu Fabra, Barcelona, Spain; Bioinformatics and Omics Data Analytics, German Cancer Research Center (DKFZ), Heidelberg, Germany; Faculty of Business Administration and Economics, Bielefeld University, Bielefeld, Germany; Department of Translational Oncology, NCT Dresden, University Hospital, Carl Gustav Carus, Technische Universität Dresden, Dresden and DKFZ, Heidelberg, Germany; German Cancer Consortium, Heidelberg, Germany; Department for Bioinformatics and Functional Genomics, Institute for Pharmacy and Molecular Biotechnology (IPMB) Heidelberg University, Heidelberg, Germany

**Keywords:** 3D cell culture, tumor cell heterogeneity, single cell analysis, gene expression deconvolution, spheroid image analysis

## Abstract

3D-culture systems have advanced cancer modeling by reflecting physiological characteristics of *in-vivo* tissues, but our understanding of functional intratumor heterogeneity including visual phenotypes and underlying gene expression is still limited. Single-cell RNA-sequencing is the method of choice to dissect transcriptional tumor cell heterogeneity in an unbiased way, but this approach is limited in correlating gene expression with contextual cellular phenotypes.

To link morphological features and gene expression in 3D-culture systems, we present ‘pheno-seq’ for integrated high-throughput imaging and transcriptomic profiling of clonal tumor spheroids. Specifically, we identify characteristic EMT expression signatures that are associated with invasive growth behavior in a 3D breast cancer model. Additionally, pheno-seq determined transcriptional programs containing lineage-specific markers that can be linked to heterogeneous proliferative capacity in a patient-derived 3D model of colorectal cancer. Finally, we provide evidence that pheno-seq identifies morphology-specific genes that are missed by scRNA-seq and inferred single-cell regulatory states without acquiring additional single cell expression profiles. We anticipate that directly linking molecular features with patho-phenotypes of cancer cells will improve the understanding of intratumor heterogeneity and consequently be useful for translational research.

## Introduction

Three-dimensional (3D) cell culture systems provide a physiologically relevant context for *in-vitro* testing, manipulation and high-throughput screening applications^1^. Mimicking the 3D-tissue environment thus holds great promise for future diagnostics^2^ and the analysis of functional differences between tumor cells in a single patient (intratumor heterogeneity)^3^, a phenomenon increasingly recognized as an essential driver of tumorigenic progression, trea™ent resistance and relapse^4^.

Single-cell 3D-culture combined with microscopy and molecular analyses appears as a key strategy for investigating cellular heterogeneity *in-vitro* as it enables analysis of clonal behavior in defined spatial and temporal conditions^5,6^. Ideally, the visual phenotype of the self-organizing multicellular complex (spheroids, organoids, etc.) reflects the characteristics of the primary tumor and consequently informs about the functional outcome of heterogeneous cancer cell states. While visual characteristics of 3D-cultures such as shape and size can be highly informative for classification of tumor subtypes and disease states^2,7^, most studies have so far focused on inter-patient differences rather than heterogeneous behavior of cells derived from a single patient^8^.

In primary samples, histopathology and associated visual observation of contextual cellular phenotypes *in situ* has been a common strategy for tumor classification and the analysis of intratumor heterogeneity for over a century^9^. However, the number of simultaneous molecular measurements is highly restricted with imaging-based methods or they require elaborated sample processing and highly complex experimental setups^10,11^. Recently developed methods for single cell RNA-seq (scRNA-seq) ^12,13^ have greatly improved the analysis of intratumor heterogeneity by enabling the unbiased detection of transcript abundances in individual cells^14–16^ but these approaches do not provide a direct link to visual cellular phenotypes since the available protocols involve dissociation of cells and loss of their multicellular context. Alternatively, laser capture microdissection (LCM) enables the isolation of cells from histological slices by laser cutting, which has already been combined with low input gene expression profiling in archived frozen tissue^17^ and 3D cell culture systems^18^. Although this strategy maintains the spatial information of isolated cells, key limitations of LCM are the low throughput and diminished RNA sample quality. To circumvent these limitations, recent studies have demonstrated the direct combination of histopathology and RNA-seq on primary samples to spatially and morphologically resolve transcriptional intratumor heterogeneity at a cellular resolution of 10-30 cells (spatial transcriptomics) ^19,20^.

Here, we present ‘pheno-seq’ to dissect morphological heterogeneity in 3D cell culture systems by directly combining clonal cell culture, imaging and transcriptomic profiling without the necessity of histological preparation. We developed an experimental and computational workflow for unbiased high-throughput pheno-seq, including automated dispensing and imaging of single spheroids in barcoded nanowells as well as an automated image processing pipeline. We demonstrate the power of pheno-seq in dissecting both cellular and molecular heterogeneity for established and patient-derived 3D-models of breast and colon cancer, respectively.

## Results

### Pheno-seq directly links spheroid morphologies to gene expression

In breast cancer, normal epithelial cells undergo a stepwise transformation from local hyperplasia to premalignant carcinoma *in-situ* and invasive carcinoma^21^. Importantly, the switch from epithelial to invasive behavior requires gene expression programs that resemble those occurring during embryogenesis and wound healing, generally described as epithelial-to-mesenchymal transition (EMT) ^22^.

Single-cell-derived spheroids of the breast cancer cell line MCF10CA^23^ show a remarkable morphological heterogeneity when cultured in 3D, with cellular phenotypes reflecting characteristics of both carcinoma *in-situ* (‘round’ phenotype) and invasive carcinoma (‘aberrant’) (Supplementary Fig. 1a and b). To enable independent analysis of cells derived from both phenotypes, we developed a workflow to isolate single spheroids from reconstituted basement membrane (Matrigel) without perturbing their phenotypic identity (Fig. 1a and b). To functionally assess the observed heterogeneity, we reseeded and cultured cells from both phenotype classes independently which validated efficient isolation and revealed a high cell state stability (Supplementary Fig. 1c).

**Figure 1.**
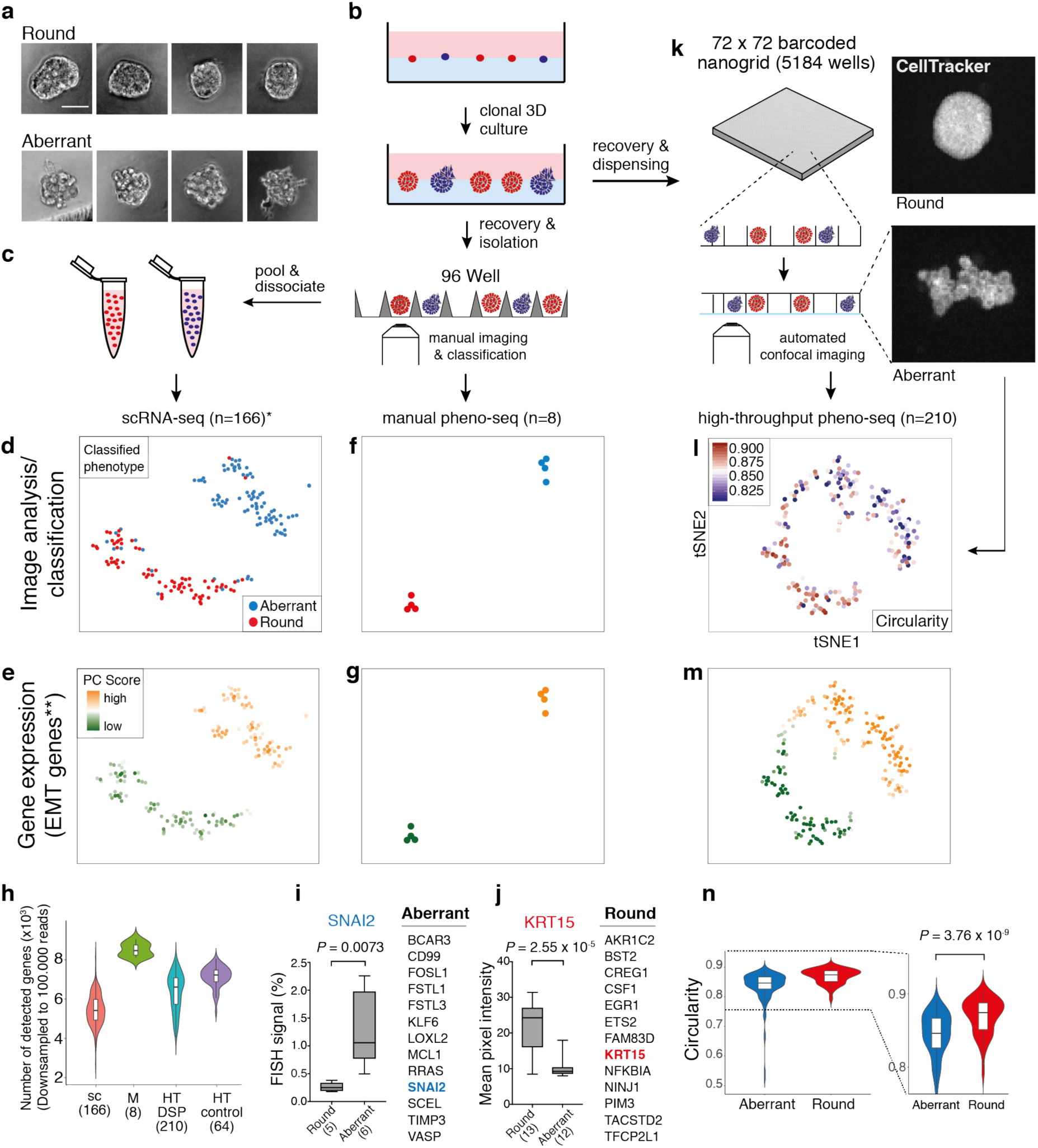
pheno-seq enables direct image correlation and complements the identification of morphology-specific gene expression. **(a)** Brightfield images of clonal spheroids (MCF10CA phenotype classes ‘round’ and ‘aberrant’) after isolation from Matrigel (scale bar 50 µm). **(b)** Workflow overview for the isolation of clonal spheroids for inference of morphology-specific gene expression. **(c)** Indirect phenotype – gene expression correlation by scRNA-seq using single cells isolated from multiple spheroids with annotated morphology phenotype. **(d)** 2D tSNE visualization^24^ of 166 scRNA-seq (*cell-cycle corrected) full-length expression profiles of cells from manually isolated round and aberrant spheroids with coloring based on manual phenotype annotation. **(e)** Same 2D tSNE visualization as shown in (d) but coloring based on PC scores for **HALLMARK_EMT gene set derived from the Molecular Signature Database (MSigDB)^25^. **(f and g)** 2D tSNE visualization of 8 full-length manual pheno-seq profiles based on manually isolated single spheroids. Same coloring as shown in (d) and (e). **(h)** Number of genes detected in downsampled scRNA-seq and pheno-seq libraries (sc: scRNA-seq; M: manual pheno-seq; HT-DSP: high-throughput pheno-seq with dithio-bis(succinimidyl) propionate fixation; HT-control: HT-pheno-seq bottom control). Numbers of samples indicated on x-axis under respective method. **(i and j)** Selected genes only identified by manual pheno-seq and not by scRNA-seq (Differential expression analysis^26^: Fold change > 1.3; adjusted *P*-value < 0.1) and validation of phenotype-specific expression for SNAI2 (aberrant) and KRT15 (round). RNA-FISH for SNAI2: Plotted values reflect the faction of pixels that exceed the background threshold per spheroid. KRT15 immunofluorescence: Plotted values reflect mean pixel intensity per classified spheroid. Box plot center-line: median; box limits: first and third quartile; whiskers: min/max values. Numbers of samples indicated on x-axis under respective phenotype class. Indicated are *P*-values from unpaired two-tailed Students t-test. **(k)** High-throughput (HT) pheno-seq workflow based on automated dispensing and confocal imaging of recovered spheroids in barcoded nanowells. **(l)** 2D tSNE visualization of 210 HT-pheno-seq 3′-end profiles with coloring based on image feature ‘circularity’. For better visualization, all circularity values below 0.8 were set to minimum in the color code scheme. **(m)** Same 2D tSNE visualization as shown in (l) with coloring based on PC scores for **HALLMARK_EMT gene set as shown in (e) and (g). **(n)** Circularity plotted per cluster (k-means clustering, k=2) as shown in (l). Violin-plot center-line: median; box limits: first and third quartile; whiskers: ±1.5 IQR. Indicated *P*-value from unpaired two-tailed Students t-test.

As a reference dataset, we first generated and deeply sequenced microfluidics-based full-length scRNA-seq libraries of cells derived from both spheroid phenotypes independently (166 cells in total, Fig. 1c, Supplementary Table 1). Notably, this strategy does not provide a direct link between spheroid morphologies and gene expression as multiple spheroids needed to be pooled to acquire a sufficiently high number of input cells.

Combined transcriptomic analysis by testing annotated and *de-novo* identified gene sets for coordinated expression variability^24^ and t-SNE visualization revealed two distinct clusters and a tight association of cells to their original phenotype class (Fig. 1d), whereas differential expression analysis^26^ identified biologically relevant expression patterns. Cells derived from aberrant spheroids are characterized by the expression of EMT related genes (Fig. 1e), including vimentin (VIM), Beta-Actin (ACTB) and fibroblast activating protein (FAP). In contrast, cells isolated from round spheroids showed higher expression of genes involved in adherence and formation of tissue structures including desmoglein 3 (DSG3) and keratin 16 (KRT16) (Supplementary Fig. 2a and 3a). In order to validate if we could accurately detect gene expression specific for invasive phenotypes, we used whole mount immunofluorescence (IF) of individual marker genes, in particular the EMT marker VIM and the cytoskeleton component ACTB (Supplementary Fig. 4a and b).

**Figure 2.**
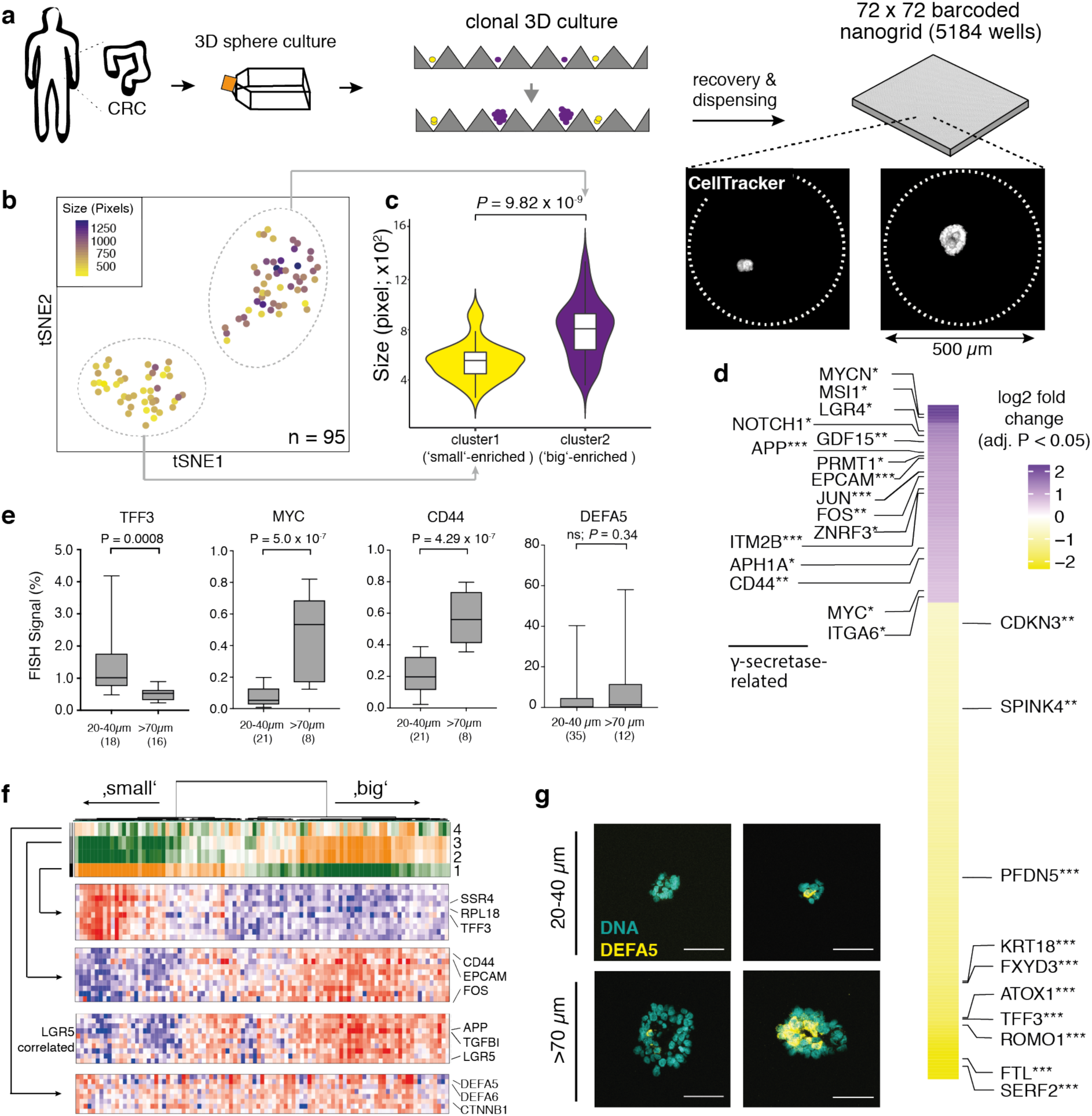
pheno-seq of a 3D model of colorectal cancer links heterogeneous proliferative phenotypes to expression signatures enriched for cell type-specific markers. **(a)** Clonal 3D-culture in inverse pyramidal shaped microwells and recovery strategy for HT-pheno-seq of patient-derived CRC spheroids isolated from a liver metastasis. Yellow and purple indicate heterogeneous subpopulations with functional differences in proliferative potential^33^. **(b)** 2D tSNE visualization of 95 HT-pheno-seq expression profiles. Coloring by sphere size (pixel). **(c)** Spheroid size plotted per cluster. Violin-plot center-line: median; box limits: first and third quartile; whiskers: ±1.5 IQR). Indicated P-value calculated from unpaired two-tailed Students t-test. **(d)** Hea™ap reflecting differential expression analysis of identified clusters in (b). Selected genes are listed beside the hea™ap; Fold change > 1.5; adjusted *P*-value < 0.05; \**P* < 0.05, \*\**P* < 0.01, \*\*\**P* < 0.001; ‘small’ cluster1: 313 differentially expressed genes; ‘big’ cluster: 130 differentially expressed genes. **(e)** Validation of pheno-seq by quantitative RNA-FISH for size-dependent differentiation marker TFF3 and stem cell markers CD44/MYC, and for size-independent Paneth-cell marker DEFA5. Plotted values reflect the pixel fraction that exceeds the background threshold per spheroid (Box plot center-line: median; box limits: first and third quartile; whiskers: min/max values; *P*-values from unpaired Students t-test, ns: non-significant. Numbers of samples n indicated on x-axis under respective class). **(f)** PAGODA RNA-seq analysis of CRC spheroid HT-pheno-seq data. Dendrogram indicates overall clustering and the rows below represent top four significant aspects of heterogeneity based on HALLMARK/GO gene sets derived from the MSigDB and on de-novo identified gene sets. High aspect scores (PC Scores) correspond to high expression of associated gene sets. Expression patterns below reflect top 10 loading genes for selected gene sets that are associated with respective aspects. Bottom: Expression pattern of top genes most highly correlated with stem cell marker LGR5 and Paneth marker DEFA5 (Pearson’s correlation) **(g)** Example images (Z-projections) for RNA-FISH staining of Paneth-cell marker DEFA5 corresponding to data shown in (e). DNA (Hoechst) counterstain visualization (Hoechst: cyan; RNA: yellow; scale bar 50 µm).

We next tested whether expression profiling of manually isolated whole spheroids (manual pheno-seq) might serve as a complementary approach to identify transcriptional differences between clonal spheroid phenotypes. Notably, this approach directly links observed heterogeneous morphologies to underlying gene expression and provides more RNA material for cDNA library preparation. Profiling of only eight spheroids by manual pheno-seq yielded a similar phenotype-specific clustering defined by high and low expression of EMT-related genes (Fig. 1f and g, Supplementary Fig. 2b). While the sample number was approximately 20 times lower (166 single-cells vs. 8 single spheroids), the gene detection rate per sample was significantly higher compared to scRNA-seq (Fig. 1h), and differential expression analysis revealed over 100 morphology-specific genes that could not be detected by scRNA-seq (Fig. 1i and j, Supplementary Fig. 3b). Although we detected more differentially expresses genes by scRNA-seq, most likely due to the much higher sample number, only pheno-seq identified the transcriptional EMT master regulator SNAI2^27^ (aberrant) and keratin 15 (KRT15, round), a basal-myoepithelial marker in the mammary gland^28^ (Fig. 1i and j, Supplementary Fig. 3c).

**Figure 3.**
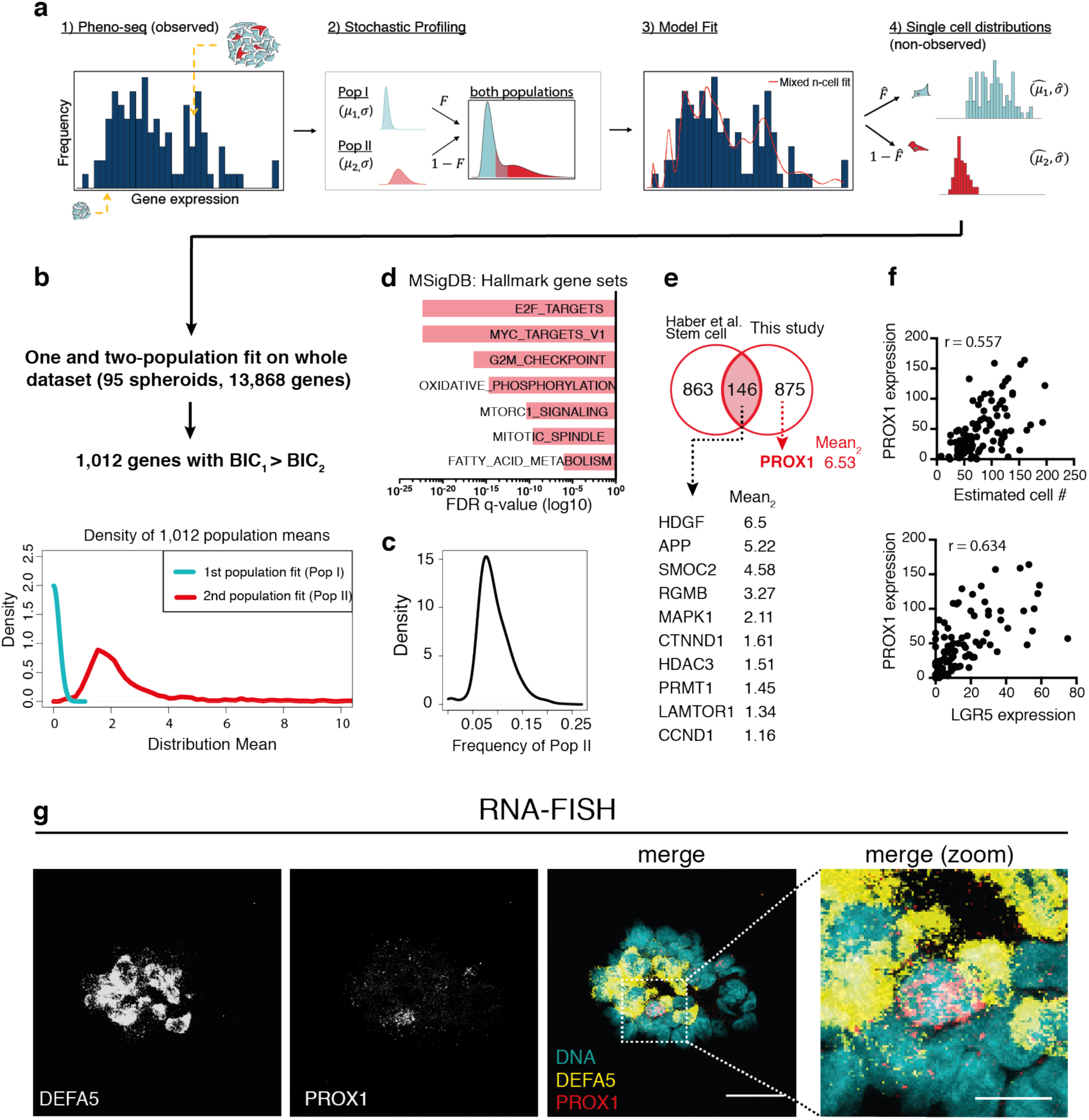
Single-cell deconvolution of CRC spheroid pheno-seq data by maximum likelihood inference identifies PROX1 as potential CRC stem cell marker. **(a)** Concept of adapted maximum likelihood approach^41^ based on estimated cell numbers and transformed pheno-seq data (n = 95): 1) Acquired and transformed pheno-seq data based on estimated cell numbers build a distribution of measurements for inference by the model. Coloring of cells in spheroids: red = stem-like; cyan = differentiated. 2) Assumptions on single cell distributions: Model of heterogeneous gene regulation in which single cells are supposed to exhibit gene expression at low (Pop I) or high (Pop II) levels with a common coefficient of variation. The four parameters of the model are the log-mean expression for each subpopulation (μ1 and μ2), the proportion of cells in the high subpopulation (*F*), and the common log-SD of expression (σ). 3) Based on the model in step 2, a likelihood function is derived that takes different numbers of cells per spheroid into account. The likelihood function is then maximized by searching through the four parameters of the model to identify those that are most likely given the experimental observations. 4) These four parameters define the inferred single cell distributions of the low and high-level populations. **(b)** 1,012 genes show an improved two-population fit compared to a one population fit (BIC: Bayesian information criterion). Densities of the means of the first (Pop I: low regulatory state) and second population (Pop II: high regulatory state) for all identified 1,012 genes. **(c)** Frequency distribution of cells with high regulatory state (Pop II) of identified 1,012 genes. **(d)** Gene set enrichment analysis for two-population genes based on Hallmark gene sets derived from the MSigDB. Bar plot showing top enriched gene sets ranked by FDR q-values. **(e)** Venn-diagram showing overlap between identified two-population genes and murine small intestinal stem cell signature from scRNA-seq study^36^. Selected genes are listed below ordered by mean for high-state population Pop II (Mean2). **(f)** Scatter plots for PROX1 expression plotted against estimated cell numbers (upper) and against expression of major intestinal stem cell marker LGR5 (lower) as well as associated Pearson’s correlation coefficients (r). **(g)** RNA-FISH co-staining of CRC spheroids for PROX1 (Atto550) and DEFA5 (Alexa488) and Hoechst counterstaining for visualization of DNA. Merged images: DNA: cyan; DEFA5 yellow; PROX1: red. Images represent Z-projections (scale bar 30 µm and 10 µm for magnified merged image).

Phenotype-specific expression of SNAI2 and KRT15 was validated by RNA-FISH and immunofluorescence (IF), respectively (Fig.1i and j, Supplementary Fig. 4c and d). We reasoned that SNAI2 could not be identified by scRNA-seq due to its low expression (Supplementary Fig. 3c), a frequent phenomenon for transcriptional regulators^29^. Although KRT15 is one of the top markers for round spheroids detected by pheno-seq, the existence of residual KRT15^+^ cells in aberrant spheroids (Supplementary Fig. 4c) seemed to mask the identification of KRT15 as phenotype-specific when single-cell profiles were analyzed. Remarkably, differential expression of KRT15 and SNAI2 could not be robustly restored from scRNA-seq data by generating synthetic pheno-seq profiles from averaged single-cell expression (Supplementary Fig. 2c and Supplementary Fig. 2d), indicating for the additional influence of dissociation bias^30^ on KRT15 mRNA abundance. In summary, pheno-seq provides the direct link between spheroid morphologies and underlying transcriptomes and complements scRNA-seq methods in identifying expression differences between heterogeneous spheroid phenotypes already with low sample numbers.

### High-throughput pheno-seq in barcoded nanowells enables combined quantitative analysis of image features and gene expression

A major limitation of both scRNA-seq and manual pheno-seq is the non-quantitative and biased selection of spheroid phenotypes based on visual inspection by eye. In addition, increasing the number of spheroids per pheno-seq experiment is necessary to comprehensively understand associations between visual phenotypes and gene expression in a particular 3D-culture model. Therefore, we developed high-throughput (HT) pheno-seq by repurposing the nanowell-based iCELL8 scRNA-seq system^31^, a platform for integrated imaging and gene expression profiling of single cells, for the processing of spheroid samples of up to 150 µm in size. Key modifications included cellular fixation^32^, altered chip setup, higher-resolution microscopy, an automated image-processing pipeline and the ‘PhenoSelect’ software for interactive analysis and selection of spheroids for sequencing (Fig. 1k, Supplementary Fig. 5, 6 and 7). These substantial technical adaptions had only minor influences on the gene detection rate, which fell in between scRNA-seq and manual pheno-seq (Fig 1h, Supplementary Table 1). MCF10CA HT-pheno-seq yielded very similar results as described, with two distinct clusters driven by expression of genes involved in EMT (VIM^+^) as well as tissue formation (KRT15^+^) but at higher throughput per experiment (n = 210) compared to manual pheno-seq (Fig 1l and m, Supplementary Fig. 8a). Both pheno-seq methods show good concordance in identifying differentially expressed genes between spheroid phenotypes (Supplementary Fig. 8b), despite unbiased capture of spheroids by HT-pheno-seq as well as differences in sample size and library structure (3′-end vs. full-length, Supplementary Table 1).

In contrast to scRNA-seq, HT-pheno-seq allows measurements of RNA abundance and image features from the same spheroid, which enabled straightforward association of genetic programs and complex visual phenotypes based on the fluorescence signal derived from a cytoplasmic dye (CellTracker Red). These included the morphology-related feature ‘circularity’, informing about (de)regulation of lobular development (Fig. 1l and n), and spheroid size, demonstrating a higher proliferative activity of epithelial-like cells (Supplementary Fig. 8c). In addition, pheno-seq linked negatively skewed pixel intensity distributions to round phenotypes (Supplementary Fig. 8d), indicative of increased cell density in round 3D phenotypes that leads to an increased proportion of high pixel intensity values derived from the cytoplasmic signal. Hence, HT-pheno-seq represents a new method that directly and quantitatively links heterogeneous spheroid phenotypes to underlying gene expression in a single experiment.

### HT-pheno-seq of a patient-derived colorectal 3D model links proliferative capacity to cell type-specific expression signatures

We next set out to assess the functional correlation between visual phenotypes and gene expression in a physiologically relevant and patient-derived 3D model originally isolated from a liver metastasis of a colorectal cancer (CRC) patient. Similar to the phenotypic heterogeneity in the MCF10CA spheroids described above, functionally distinct subpopulations in 3D-cultures of CRC patients have previously been identified^33^. The reported heterogeneity in proliferative potential seems to be largely independent of mutational subclone diversity^34^, thereby supporting the presence of a differentiation-like hierarchy in CRC^35^. As reseeding of cells from different classes of spheroid sizes (20-40 µm and >70 µm) revealed significant differences in spheroid forming capacity (Supplementary Fig. 9a and b), we hypothesized that specific stem- and differentiation-related transcriptional signatures should underlie these heterogenous proliferative phenotypes. To investigate this hypothesis, we performed HT-pheno-seq based on clonal CRC spheroids cultured in an inverse pyramidal-shaped microwell setup (Fig. 2a; Supplementary Fig. 9c).

Analysis of relative gene expression differences between 95 HT-pheno-seq profiles and t-SNE visualization confirmed two transcriptionally distinct clusters (Fig. 2b). Image analysis of the respective spheroids revealed a strong difference in spheroid size composition between both clusters (Fig. 2c) that does not influence library complexity (Supplementary Fig. 10b). Differential expression analysis showed that the first cluster (‘small’ phenotype) is enriched for genes involved in ribosomal activity (GO_RIBOSOME, FDR q-value 2.41×10^-45^) as well as intestinal secretory lineage markers, including Trefoil Factor 3 (TFF3), KRT18 and SPINK4^36^ (Fig. 2d). In contrast, the second cluster (‘big’ phenotype) is characterized by the expression of genes previously described to be involved in (i) stem cell maintenance (including CD44, MYC, NOTCH1, APP, MSI1 and ITGA6) ^36,37^, (ii) the formation of cell-cell junctions (including EPCAM, CLDN4, CDH1) and (iii) WNT signaling (ZNRF3, LGR4, JUN) (Fig. 2d). The pattern of this signature showed a high overlap with genes correlated with the major intestinal stem cell marker LGR5, including CD44, APP and SMOC2 (Fig. 2f, Supplementary Fig. 10a). We validated sphere size-dependent expression for selected lineage-specific markers by quantitative RNA-FISH (Fig. 2e, Supplementary Fig. 11a-c).

In the cluster enriched for big spheres, we identified several genes related to the γ-secretase machinery (Fig. 2d), a key component of the NOTCH pathway and target of new therapies aiming to disrupt cancer stem cell signaling^38^. Importantly, selective targeting of the γ-secretase by a small molecule inhibitor in concentration ranges that have been shown to force colonic stem cells into differentiation^39^ showed a inhibitory effect on spheroid growth (Supplementary Fig. 11d). This finding suggests a similar signaling dependency of the normal and transformed intestinal stem cell niche and shows the potential of pheno-seq to identify relevant signaling components required for cellular proliferation.

Moreover, we determined an expression signature primarily driven by the expression of Paneth cell markers DEFA5 and DEFA6 that is independent of the size-related clusters shown above (Fig. 2f, Supplementary Fig. 10a). Paneth cells represent a post-mitotic secretory subpopulation at the bottom of intestinal crypts that serves as niche for LGR5^+^ stem cells^39,40^. In line with pheno-seq results, we validated high-expressing DEFA5^+^ cells as rare subpopulation with spheroid size-independent relative expression by RNA-FISH (Fig. 2e and g, Supplementary Fig. 10c). It has been shown previously that the percentage of functionally different subpopulations remains stable over several rounds of replating, suggesting that the composition of cells in continuously growing spheroids remains stable^33^. Consequently, for a cellular subtype with limited proliferative potential within the putative CRC differentiation hierarchy, we would have expected a similar association of relative expression and size for DEFA5^+^ cells as observed for the TFF3^+^ secretory signature above. However, as we could detect several big spheres with very high numbers of DEFA5^+^ cells, we suggest that Paneth-like cells exhibit a heterogeneous proliferative phenotype (high- and low-cycling) that might relate to the delayed-contributing subpopulation in CRC previously described^33^. Thus, pheno-seq is able to directly assign heterogeneous proliferative phenotypes to expression signatures enriched for specific intestinal cell-type markers.

### Single-cell deconvolution by combining image analysis and maximum likelihood inference

The pheno-seq method enables the direct association of spheroid morphologies and gene expression. However, this ability comes at the cost of lower cellular resolution. The gene expression signatures identified from CRC spheroids inform about general phenotype-specific expression and trends in subtype composition but might derive from multiple cellular subtypes present within the same spheroids. While these results are highly valuable for understanding growth behavior in clonal cell culture systems, obtaining ‘real’ single-cell information from pheno-seq data would be of high relevance to distinguish between genes that are generally associated with spheroid phenotypes and those who are robustly expressed in a single-cell subpopulation. Therefore, we aimed to computationally infer single-cell regulatory states by deconvolution of gene expression data using both image analysis and a maximum likelihood inference approach.

First, we generated a 3D high-resolution imaging reference dataset by light-sheet microscopy from spheroids of different sizes, which we used to determine the relationship of spheroid size and nuclei counts to estimate cell numbers from CRC spheroid pheno-seq imaging data (Supplementary Fig. 12a). As the original pheno-seq data exhibited a low correlation between library complexity and estimated cell numbers, we downsampled the data to achieve a constant number of mRNA counts per estimated single cell content (Supplementary Fig. 12b). As expected, this transformation introduces a positive overall shift of correlations between gene expression and cell numbers (Supplementary Fig. 12c), which can be mainly explained by housekeeping genes with a constant number of mRNA molecules per cell (Supplementary Fig. 13a). However, heterogeneously expressed genes such as the differentiation markers TFF3 and DEFA5 do not exhibit any correlation with cell numbers (Supplementary Fig. 13b and c), validating our normalization approach.

To identify genes whose expression was likely to be informative for heterogeneous single-cell regulatory states, we used a maximum likelihood inference approach initially developed to deconvolve cell-to-cell heterogeneities from random 10-cell samples41 (Fig. 3a). The adapted algorithm uses the estimated cell numbers per spheroid to fit two log-normal distributions (LN-LN model) to given ‘mixed-n’ datasets in order to identify genes with bimodal expression pattern at the single-cell level (Stochastic Profiling, see Methods). Importantly, this approach unbiasedly pinpoints genes that show a heterogeneous and robust expression within spheroids at the single-cell level, instead of comparing gene expression between spheroids.

Whilst the deconvolution technique assumes that cellular subtypes are identically distributed across samples, pheno-seq is based on clonal spheroids whose cell number, subtype composition and expression profile is dependent on the state of the founding cell. Based on the cancer stem cell model and the CRC differentiation hierarchy confirmed above, we assume that continuously growing spheroids (‘big’ phenotype) harbor all cellular subtypes present in this system, including stem-like cells, whereas small spheroids with limited proliferative capacity and low cell numbers are more homogeneous and contain only differentiated subtypes. Thus, inferred regulatory states should be enriched for genes specific for the stem-like compar™ent, as these represent a major source of heterogeneity across all spheroids at the single-cell level.

Deconvolution of the entire CRC pheno-seq dataset revealed 1,012 genes that show an improved two-population fit compared to a one-population fit, assessed by the Bayesian information criterion (BIC) to calculate the quality of the fit relative to the number of inferred parameters (Fig. 3b). Most of the fits resulted in a highly-expressing cellular fraction of 5 – 15% (Fig. 3c) thereby matching the proportion of cells with spheroid forming capacity in this model^33^. Interestingly, the positive shift of correlations between gene expression and cell numbers (before and after downsampling) is much more pronounced in the set of two-population genes compared to the set of non-two-population genes (Supplementary Fig. 13d), suggesting that many of the inferred two-population genes are involved in proliferative potential. Indeed, gene set enrichment analysis reveals a high proportion of MYC targets as well as genes involved in the regulation of cell growth and proliferation (Fig. 3d). In addition, high enrichment of genes involved in oxidative phosphorylation indicates for heterogeneous mitochondrial activity at the single-cell level, a phenomenon recently described for intestinal stem cells and niche-forming Paneth cells in the small intestine^42^. Strikingly, a high number of identified genes are overlapping with a recently identified stem cell signature of the small intestine revealed by massively parallel scRNA-seq^36^, including SMOC2, APP, PRMT1, RGMB, MAPK1 and CTNND1, respectively (Fig 3e).

Here, we identified the transcriptional regulator PROX1 as gene with a high population (Pop II) mean (Fig. 3e) that is strongly correlated with cell numbers and with expression of the major stem cell marker LGR5 (Fig. 3f). In addition, PROX1 top correlated genes exhibit a strong overlap with the signature defining big spheres when relative gene expression differences between spheroids were analyzed (Supplementary Fig. 10a). In the normal intestinal epithelium, PROX1 is expressed in the enteroendocrine lineage^43^. However, two studies based on mouse tumor models suggest a role for PROX1 in cancer stem cell maintenance and metastatic outgrowth^44,45^. In line with these observations, we validated PROX1^+^ cells by RNA-FISH as a rare subpopulation in a patient-derived human tumor model (Fig. 3g). Furthermore, PROX1+ cells are framed by DEFA5^+^-positive Paneth-like cells, suggesting a similar niche dependency for normal stem cells and CRC stem-like cells at distant sites of neoplasia. Taken together, gene expression deconvolution of pheno-seq data provides information about gene expression patterns at the single cell level even without acquiring additional single cell expression profiles.

## Discussion

As applications of 3D-cultures are emerging even into clinical settings^2^, there is increasing need to directly combine unbiased gene expression profiling with *in situ* imaging to understand heterogeneous oncogenic phenotypes, similar to spatial transcriptomics in primary tumor samples^19,20^.

In this study, we present pheno-seq as new and complementary approach that directly combines high-throughput imaging and next generation sequencing to explain heterogeneous morphologies of clonal tumor spheroids. At the same time, pheno-seq bridges the gap between single-cell and bulk expression profiling and complements scRNA-seq in identifying heterogeneous gene expression. In principle, pheno-seq can be applied to any 3D-culture system given that the phenotypic identity is maintained upon spheroid isolation. For example, morphologies of organoids derived from patients with pancreatic ductal adenocarcinoma can be linked to the state of malignant transformation and prognosis^46^. We expect that this combination of functional single cell growth assay with combined image and gene expression profiling will be widely applied in cancer biology, ranging from primary to circulating tumor cells (CTCs^47^).

Importantly, we show that pheno-seq is able to link cell type-specific genes to heterogeneous growth phenotypes even in highly complex cell culture systems. In addition, we show that deconvolution by maximum likelihood inference provides an additional layer of information by revealing single-cell regulatory states that are likely to be associated with a distinct stem-like population, thereby further supporting a differentiation-like hierarchy in CRC. Based on our results, future studies should shed light on additional functional characteristics and dependencies of the stem-like compar™ent, the implication of the heterogeneous growth phenotype of Paneth-like cells, potential cancer cell plasticity and the impact of subtype-specific metabolic preferences. Similar to a recent study integrating scRNA-seq and spatial transcriptomics^48^, complementary single-cell-derived information might be added to further deconvolve cell type compositions in pheno-seq expression profiles.

We envision pheno-seq to become even more powerful with increasing resolution and content of imaging, employing enhanced 3D-image acquisition, integrated staining by IF or live-dyes, and time-lapse microscopy, respectively. Pheno-seq can also be easily extended to other low-input, next-generation sequencing modalities such as chromatin accessibility sequencing^49^. Furthermore, pheno-seq might be applied to pooled-screening approaches^50^ or to resolve transcriptional changes that are associated with morphological transitions in non-synchronized developmental processes. Thus, pheno-seq will widely impact the way how we study functional heterogeneity in a variety of biological and clinical applications.

## Acknowledgments

We thank David Ibberson (CellNetworks Deep Sequencing Core Facility, Heidelberg University) for NGS services, Daniel Liber and Marizela Kulisic (TakaraBio) for technical support for the iCELL8 system, Henrik Kaessmann (ZMBH, Heidelberg University) for support and helpful discussions regarding the iCELL8 system and single cell analysis, Lorenz Maier (Theoretical Bioinformatics, DKFZ) for help with KNIME, Katharina Jechow (Theoretical Bioinformatics, DKFZ) for technical laboratory support, Claudia Ernst and Niels Grabe (Hamamatsu TIGA Center, Heidelberg University) for help with histological preparation, Naveed Ishaque (Theoretical Bioinformatics, DKFZ) for assistance in RNA-seq data analysis and Dominik Niopek, Luca Tosti, Julia Neugebauer, Teresa Krieger and Lorenz Chua (Theoretical Bioinformatics, DKFZ) for critically revising the manuscript. Primary human colon cancer samples were obtained from Heidelberg University Hospital in accordance with the declaration of Helsinki. Informed consent on tissue collection was received from each patient, as approved by the University Ethics Review Board. ST is recipient of the stipend for the PhD program of the Helmholtz International Graduate School for Cancer Research (DKFZ, Heidelberg). This study was supported by the Helmholtz International Graduate School for Cancer Research, the iMed Program (Helmholtz Association), the BMBF-funded Heidelberg Center for Human Bioinformatics (HD-HuB) within the German Network for Bioinformatics Infrastructure (de.NBI) (#031A537A, #031A537C), the DFG (SFB873), the EU framework programme Horizon2020 (TRANSCAN-2 ERA-NET), the German Cancer Aid (Colon-Resist-Net), NCT3.0_2015.4 TransOnco. and NCT3.0_2015.54 DysregPT, the German Research Foundation (DFG) within the Collaborative Research Centre 1243, Subproject A17, the BMBF (grant # 01ZX1711A) and the Helmholtz Association (Incubator grant sparse2big, grant # ZT-I-0007). DKFZ-HIPO provided technical support and funding through Grant No. HIPO-H012.

## Author contributions

SMT and CC conceived the study, SMT, CC, HG and RE designed experiments; SMT performed 3D cell culture experiments, IF/RNA-FISH stainings and iCELL8 sample and library preparation; SMT and FP performed confocal microscopy; BE performed light-sheet microscopy; FP developed the HT-pheno-seq imaging protocol, the image processing pipeline and PhenoSelect; FP and MW performed image analysis; JPM and SMT performed Fluidigm C1 experiments and JPM generated sequencing libraries; JP, LA, ZG, SMT and SS analyzed RNA-seq data; LA, CF and FJT developed and applied the adapted maximum likelihood inference deconvolution approach for pheno-seq data; CB and HG generated and characterized the colon spheroid cultures and contributed experimental and clinical expertise; KR, MS, MG and IG provided advice on single-cell sequencing experiments and analysis. MG, IG contributed NGS expertise and sample processing. SMT and CC wrote the manuscript. All authors revised and approved the manuscript.

## Competing financial interests

The authors declare no competing financial interests

## Methods

### Breast cancer model MCF10CA Cell culture

For 2D cell culture, the cell line MCF10CA1d clone 1 (acquired from The Barbara Ann Karmanos Cancer Institute), a transformed derivative of the MCF10A 3D-culture model for acinar morphogenesis of the mammary gland, was routinely passaged in 25 cm2 culture flasks (greiner bio-one). Cells were cultured in growth medium consisting of DMEM/F12 medium supplemented with 5% horse serum, 10 µg/ml Insulin (Life Technologies), 20 ng/ml EGF, 0.5 mg/ml hydrocortisone and 100 ng/ml Cholera toxin (Sigma). Cells were passaged at approximately 80% confluency with 0.05% Trypsin (Life Technologies). The cell line was authenticated using a Multiplex human Cell line Authentication test (http://www.multiplexion.de/).

For 3D ‘on top’ assays, cells were cultured in assay medium (growth medium with only 2% horse serum and 5 ng/ml EGF) in 24-well cell culture plates (greiner bio-one). As a basement membrane surrogate, a bed of laminin-rich hydrogel (Matrigel^®^, Corning) was generated by adding 70 µm cold Matrigel into the center of pre-wetted wells. The Matrigel bed was then dried for 20 min at 37°C. For single-cell seeding, 2D cultures were dissociated into single-cell suspensions, washed once in assay medium, passed through a 35 µm strainer and counted by a LUNA™ automated cell counter (Logos Biosystems). Subsequently, 4000 cells were seeded per well in 400 µl assay medium with 5% Matrigel by adding cell suspensions in a 45° angle to the wall of the well, which resulted in uniform distribution of single-cells throughout the well. Medium was replaced every 3 days and cells cultured for up to 12 days. All scRNA-seq and pheno-seq experiments were carried out after 5 days in 3D-culture.

## Spheroid recovery from hydrogel

After MCF10CA cells were cultured in 3D for 5 days, medium was removed from wells and 500 µl filtered and pre-warmed Dispase (Sigma) was added. The hydrogel-matrix was detached from wells by scratching over the well bottom with a 1000 µl pipette tip and the whole Dispase-Matrigel suspension was carefully resuspended five times. Afterwards, spheroids were incubated at 37 ° C for 7 min. Spheroids were then transferred to a 15 ml falcon and 5 ml assay medium was added and resuspended slowly with a 5 ml pipette. Subsequently, spheroids were spun down(300 g, 3 min) and resuspended in DMEM (Life Technologies). We do not recommend using PBS due to perturbation of spheroid morphology. In general, this procedure resulted in approximately 2000 isolated spheroids per well.

### Spheroid isolation and dissociation to single-cell suspensions

In order to isolate and classify individual MCF10CA spheroids prior to dissociation, suspensions were diluted to 100 spheroids per ml in assay medium and distributed into GravityTRAP™ ultra-low attachment 96-well plates (PerkinElmer, 10 µl per well). Plates were centrifuged for 2 min at 250 g. The V-shaped wells with 1 mm diameter flat-bottom for efficient classification (round vs. aberrant) of spheroids with 10x or 20x objectives of an inverted brightfield microscope. After 30-40 spheroids had been isolated and identified for each class, 50 µl Accumax (Sigma) was added to each well followed by an incubation of 10 min at 37 ° C. To stimulate dissociation, shear forces were applied by resuspending wells of one class with a 200 µl pipette without changing the tip. After a second incubation of 5 min at 37 ° C, wells of one class were pooled in 1.5 ml microcentrifuge tubes, spun down at 300 g for 3 min and resuspended in either assay medium or DMEM/F12.

### Reseeding assay

For independent reseeding of round and aberrant 3D phenotypes, 30-40 spheroids were isolated, dissociated and pooled as described above. A 10 µl Matrigel bed was prepared during dissociation in 15µ angiogenesis slides (Ibidi). After centrifugation, cells were resuspended in 50 µl assay medium (+ 5% Matrigel) and added to pre-treated angiogenesis slides. Medium was replaced every 3 days and cells were cultured for up to 6 days.

### Single-cell capture, mRNA library preparation and sequencing

For single-cell RNA sequencing experiments, spheroids were dissociated as described above and resuspended in DMEM/F12 medium. Capture, full-length cDNA synthesis and amplification was performed on the C1 Single-Cell Auto Prep IFC (Fluidigm). Cells at a concentration of 350 cells/µl were mixed with C1 Cell Suspension Reagent (Fluidigm) at a ratio of 4:1 immediately before loading on the IFC. Single-cell capture was assessed with an inverted brightfield microscope. Workflow and reagents for single-cell RNA extraction, reverse transcription (RT) and mRNA amplification (18 cycles) were used as described in the SMARTer Ultra Low RNA Kit (for Fluidigm C1). Sequencing libraries were generated with the Nextera XT kit (Illumina) according to an adapted Fluidigm protocol. Concentration and quality of cDNA and sequencing libraries was assessed by a fluorometer (Qubit) and by electrophoresis (Agilent Bioanalyzer high sensitivity DNA chips). Libraries of up to 24 cells were pooled and sequenced as 1 × 50-bp reads on an Illumina HiSeq 2000 machine.

### Manual pheno-seq workflow, library preparation and sequencing

For manual pheno-seq, suspensions were diluted to 500 spheroids per ml in DMEM and 2 µl was carefully dispensed to the wall of the well of GravityTRAP™ 96-well plates followed by vertical tapping of the plate. Wells with single spheroids were then microscopically classified. For RNA extraction with the Arcturus PicoPure kit (ThermoFisher), 50 µl extraction buffer was directly added to 96-wells, incubated for 2 min at RT and then transferred to 1.5 ml LoBind microcentrifuge tubes (Eppendorf). RNA was isolated as described in the PicoPure Kit (Appendix B and Section 4B.2) including on-column DNase digestion (Appendix A, RNase-Free DNAse Set, Qiagen). RNA was eluted in Nuclease-free water (∼10 µl) and used as input for full-length cDNA synthesis and amplification (16 cycles) by the SMART-Seq^®^ v4 Ultra Low Input RNA Kit for sequencing (TakaraBio). Sequencing libraries were generated with the Nextera XT kit (Illumina) as described in the SMART-Seq^®^ v4 protocol. Concentration and quality of cDNA and sequencing libraries was assessed by a fluorometer (Qubit) and by electrophoresis (Agilent Bioanalyzer high sensitivity DNA chips). Ten libraries were pooled and sequenced as 1 × 50-bp reads on an Illumina HiSeq 2000 machine.

### High-throughput pheno-seq workflow, library preparation and sequencing

For high-throughput (HT-)pheno-seq, we adapted and improved the nanowell-based Wafergen iCELL8 scRNA-seq system, that integrates imaging and gene expression profiling of big samples of up to 100 µm^51^. First, spheroids were stained 3 hours with 10 µM CellTracker™ Red CMTPX dye and 1 µg/ml Hoechst 33258 (ThermoFisher). Afterwards, spheroids of six wells were recovered as described above and washed once with 7 ml DMEM (Life Technologies). Only three wells were pooled per 15 ml falcon tube for centrifugation. The reversible cross-linker dithio-bis(succinimidyl propionate) (DSP) was prepared for cellular fixation as previously described^32^ and directly filtered through a 10 µm strainer (PluriSelect). Spheroids were resuspended in 400 µl DSP and incubated for 30 min at room temperature. After fixation, spheroids were washed two times with cold PBS (centrifugation at 650 g and 500 g, 3 min, 4°C) and then resuspended in 650 µl cold PBS with 1x second diluent (for iCELL8 ™) and 0.4 U/µl recombinant RNase Inhibitor (TakaraBio). Spheroids were dispensed into a barcoded 5184-nanowell chip with the iCELL8 Single-Cell System (TakaraBio) as described in the Rapid Development Protocol (in-chip RT-PCR amplification). As a control, we first dispensed, imaged and processed one chip without cellular fixation using the default settings, the standard microscope and provided CellSelect™ software.

For improved HT-pheno-seq we applied the following modifications: Between the three dispensing intervals, wells in the 384-well source plate were stirred with a 200 µl pipette tip just before intake of suspensions with the dispensing heads in order to minimize settling of spheroids and to enable even distribution in nanowells. Similar to the standard single-cell protocol, the iCELL8 chip was tightly sealed with a strongly adhesive imaging foil (TakaraBio). Instead of spinning cells to the bottom, spheroids were centrifuged upside-down to the foil (700 g, 5 min, 4°C) in order to reduce the working distance and to avoid light reflections deep inside the well during imaging. To further enhance imaging resolution, we used an inverted confocal laser-scanning microscope (Leica SP8) with a 10x objective (2×2 wells per field of view) instead of the standard and system-integrated fluorescence wide-field microscope with 4x objective (6×6 wells per field of view). Afterwards, spheroids were centrifuged to the bottom (700 g, 5 min, 4°C) and chips were frozen at −80°C. The PhenoSelect software pipeline was used for image processing as well as spheroid detection and interactive selection (for more detailed description of microscopy, image pre-processing and PhenoSelect see below). A ‘filter file’ generated by PhenoSelect was used to dispense reagents only in selected nanowells as described in the Rapid Development Protocol (TakaraBio), with the exception that we adjusted the amount of Triton-×100 to a final well concentration of 1% for spheroids lysis. The timing of spheroid recovery and consequently the maximum spheroid size (that correlates with the number of cells per spheroid/well) should not exceed 100 µm as this might negatively influence RT efficiency. In addition, lysis reagents, concentration and duration might have to be adjusted for different culture models.

After in-chip reverse transcription and cDNA amplification (18 cycles), barcoded cDNA was pooled and processed to 3′-end sequencing libraries by the Nextera XT kit (Illumina) with specific adaptions described in the Rapid Development Protocol. Concentration and quality of cDNA and sequencing libraries was assessed by a fluorometer (Qubit) and by electrophoresis (Agilent Bioanalyzer high sensitivity DNA chips). Improved HT-pheno-seq paired-end iCELL8 libraries (21 + 70) were sequenced on an Illumina NextSeq 500 machine in high-output mode. The ‘bottom control’ chip without improved imaging was sequenced on a HiSeq 2000 machine with similar settings. However, this control was only used to assess library quality and not for further downstream analysis.

### Colon TICs spheroids

#### Cell culture

Primary patient-derived colon tumor spheroid cultures were established as described previously^33^. Primary human colon cancer samples were obtained from Heidelberg University Hospital in accordance with the declaration of Helsinki. Informed consent on tissue collection was received from each patient, as approved by the University Ethics Review Board. The culture used in this study was derived from a liver metastasis. Cells were cultured in 75 cm^2^ ultra-low attachment flasks in advanced D-MEM/F-12 medium supplemented with Glucose (0.6%), 2 mM L-glutamine (Life Technologies), 4 μg/ml heparin, 5 mM HEPES, 4 mg/ml BSA (Sigma), 10 ng/ml FGF basic and 20 ng/ml EGF (R&D Systems). Growth factors were added every 4 days and medium was exchanged every 4-8 days. For dissociation to single-cell suspensions, spheroid cultures were centrifuged for 5 min at 900 rpm and resuspended in 2-4 ml 0.25% Trypsin (Life Technologies). To stimulate dissociation, shear forces were applied with a 1000 µl pipette every 5 min for 20 min in total. Subsequently, 4-8 ml stop solution (PBS with 20% heat inactivated and sterile filtered fetal bovine serum, Life Technologies) was added and cells were centrifuged for 5 min at 900 rpm. For passaging, cells were then resuspended in medium, passed through a 40 µm strainer and counted.

#### Reseeding assay

To isolate, dissociate and reseed cells from big (70-100 µm) and small (20-40µm) spheroids independently, we cultured colon spheroids for 10 days and performed a stepwise size exclusion by (reverse-) filtering with standard 100 µm, 70 µm, 40 µm and 20 µm cell strainers, respectively. Spheroids were dissociated to single-cell suspension as described above but passed through a 15 µm cell strainer and counted. Afterwards, 50,000 cells were seeded in 60 mm Ultra Low Attachment Culture Dishes (Corning). Growth factors were added every 4 days and cells cultured for 10 days. Culture dishes were shaken every day to avoid clustering of spheroids.

#### Inhibitor assay

The γ-secretase inhibitor PF-03084014 (Sigma) was dissolved in sterile and distilled water to a stock concentration of 1 mM. To assess the influence of the inhibitor on spheroid growth, cells were dissociated as described above and 20,000 cells were seeded in 24-well ultra-low attachment plates (Corning) in the presence of the inhibitor at final concentrations of 5, 10 or 20 µM. In addition, we included a solvent control for the maximum amount of added water (20 µl). Growth factors were added every 4 days and cells cultured for 10 days.

#### Single-cell culture and HT-pheno-seq of colon tumor spheroids

For single-cell cultures of colon tumor cells, spheroids were dissociated to single-cell suspensions, passed through a 15 µm cell strainer and counted. Cells were cultured in Aggrewell^™^400 6-Well plates (StemCell Technologies) in which each well contains a standardized array of around 7000 inverse pyramidal shaped microwells with a size of 400 µm. For seeding, wells were pre-treated according to the manufacturer’s instructions, washed once with PBS and once with medium. Subsequently, 3500 cells in 3 ml medium were added in a 45o angle to the wall of the well, which resulted in uniform distribution of single-cells in microwells after settling. Growth factors were added every 4 days and cells were cultured for 10 days, resulting in 300-400 spheroids (>20 µm) per 6-well. Spheroids from 4-6 plates (24-36 wells, 168,000-252,000 microwells) were harvested, pooled and washed once with FluoroBrite DMEM (Life Technologies, 900 rpm for 5 min).

HT-pheno-seq was performed as described for MCF10CA spheroids above, but with following modifications: In contrast to MCF10CA spheroids, colon spheroids did not require DSP fixation because spheroid recovery does not involve contact loss from reconstituted basement membrane (Matrigel). To minimize disassembly of spheroids during processing, cells were resuspended and dispensed in FluoroBrite DMEM instead of PBS.

### Microscopy and image analysis

#### Image processing and analysis

Generally, acquired microscopy images were processed and analyzed using KNIME Image Processing (https://www.knime.com/community/image-processing, Version 3.2.1), ImageJ (https://imagej.nih.gov/ij/), R (Version 3.3.1)/R studio (https://www.rstudio.com/) and/or Graph Pad Prism 7 (https://www.graphpad.com /scientific-software/prism/). Generally, the ggplot2 package implemented in R and Graph Pad Prism 7 were used for data visualization and the PhenoSelect webtool design is based on the shiny package (https://shiny.rstudio.com). More detailed information on image analysis can be found in the supplementary information file and in associated KNIME workflows deposited in the pheno-seq github repository (https://github.com/eilslabs/pheno-seq).

#### Assessing single-cell seeding efficiency

Wells with Hoechst 33258 (1 µg/ml) and CellTracker Red CMPTX (10 µM)-stained single-cells wells were imaged with a 10x/0.30 air objective (Leica HC PL FLUOTAR) of a confocal laser-scanning microscope (Leica SP8) one hour after seeding. MCF10CA (24-well) and colon tumor cell (6-well AggreWell400) images (three independent wells) were analyzed with custom made KNIME image analysis workflows to count seeded cell singlet and doublets.

#### Reseeding assay

For both MCF10CA and CRC 3D cultures, images were acquired on a Zeiss LSM780 Axio Observer confocal microscope equipped with a 10x/0.3 air objective (Zeiss EC PLAN-NEOFLUAR) in brightfield.

For quantification of ‘round’ and ‘aberrant’ phenotypes in MCF10CA 3D-cultures we used the random-forest based machine learning software ilastik^52^. A training dataset for the ‘Pixel classification’ option was first generated on randomly seeded and cultured spheroids, whereas classification was based on images derived from independently reseeded cells from round and aberrant 3D phenotypes. Classification was performed in a custom KNIME workflow by applying the trained model to each image.

For reseeded colon tumor cells derived from defined size classes ‘big’ (70-100 µm) and ‘small’ (20-40 µm), 8×8 images per well of the grown spheroids were automatically acquired in 6-well plates (Greiner) using a custom Zeiss VBA macro. All images were analyzed using a custom KNIME workflow to measure spheroid counts per condition.

#### γ-secretase inhibitor assay

Treated and cultured (10 days) spheroids were stained with CellTracker Red CMPTX (10 µM) for 3 hours prior to imaging with a Leica SP8 confocal laser-scanning microscope equipped with a 10x/0.30 air objective (Leica HC PL FLUOTAR). 5×5 images per well (10 Z-stacks per position) were acquired automatically using the TileScan option that directly stitches acquired images to one final composite image specifically for each well. Stiched CellTracker Red images were analyzed using a custom KNIME workflow to quantify the average spheroid size per condition.

#### HT-pheno-Seq microscopy, image processing and PhenoSelect

For inverted imaging, 5184-nanowell chips were fixed on a metallic Chip Spinner (TakaraBio) with adhesive tape and placed into a standard plate holder. All wells were imaged upside-down automatically using an inverted Leica SP8 confocal microscope system. We used a 10x/0.30 air objective (Leica HC PL FLUOTAR) but images were acquired with 0.9x digital zoom to span 4 wells per field of view. Excitation was set to 405 and 552 nm and emission filter were set to receive signals between 415 – 485 nm (Hoechst) and 555 – 625 nm (CellTracker Red), respectively. Laser intensity and gain were adapted for every experiment, but the pinhole was set to 5.0 Airy Units permanently. The ‘predictive focus’ option was used to extrapolate the correct focus position for each well. One image contained 512×512 pixels, with 2.53 μm pixel size. A pre-defined HCS A template of the LAS X microscope software (Leica) was used for the grid design matching the chip dimensions. Scanning of one chip with these settings took approximately 30 minutes, resulting in 2 x 1296 images. The first part of the PhenoSelect image analysis workflow (KNIME/ImageJ) was used for assigning images to their correct well positions, image cropping, spheroid detection and segmentation as well as feature extraction and quantification.

The second part (PhenoSelect, Supplementary Figure 4 and 10) was used for interactive analysis and final selection of wells with a web-based shiny app: The saved .csv file containing spheroid statistics was automatically handled by a custom R script and embedded together with the images in an interactive R/shiny application (PhenoSelect). This allows manual browsing through the acquired images and visualization of spheroids together with their respective image feature statistics. Moreover, the application allowed for visual inspection of the given image features over the whole population, allowing the identification of specific subtypes, e.g. by a particular shape or size. To characterize absolute spheroid sizes, the respective major axis length value (in pixels) was multiplied with the physical length of a pixel in the segmented object. Subtypes of spheroids can be selected by applying different sets of thresholds (e.g. size, circularity) and individual wells can be discarded if necessary (e.g. due to imaging artefacts). The list of selected wells can be saved at any time and also reloaded to proceed with selection at a later time-point. Furthermore, comments can be added to individual wells. Control wells can be selected individually. Once the desired number of wells to be sequenced had been selected, the application generated the ‘filter file’, which is then used to program the iCELL8 dispenser software. In addition, a ‘well-list file’ was generated that contained well-barcode assignments for demultiplexing as well as calculated image features for selected spheroids. Finally, we implemented plotting of image features on pre-computed t-SNE maps based on gene expression generated by PAGODA (see below). After sequencing of selected spheroids, this tab enabled integrative analysis for direct association of functional visual phenotypes to transcriptomic heterogeneity.

#### Leakage test

Due to the additional centrifugation step to the foil, we assessed potential leakage by dispensing a highly fluorescent solution of PBS + 1 µg/ml fluorescein sodium salt (Sigma) into one half of the nanowells and dispensed only PBS into the other half and into control wells. A dispensing pattern was chosen to generate a maximum number of borders between nanowells filled with fluorescein and those only filled with PBS. Subsequently, the chip was processed and imaged as described above but with laser and filter sets matching the fluorescent properties of fluorescein (λ ^ex^ 460 nm; λ ^em^ 515 nm). The acquired images were processed for well assignment, segmentation and cropping by the HT-pheno-seq pre-processing workflow and average fluorescence intensity was measured for every well independently. Average fluorescence intensity values ranging from 0 to 255 (8 bit) were color coded and plotted onto a 72×72 grid resembling the iCELL8 chip layout using a custom R script.

#### Antibody staining for immunofluorescence

MCF10CA cells cultured in 3D were prepared for immunofluorescence staining as described previously^53^. Briefly, cells were fixed in 24-wells with 2% Formaldehyde solution (Methanol-free, ThermoFisher) for 20 min at RT and washed twice with PBS. Cells were permeabilized with PBS + 0.5% TritonX-100 (Sigma) for 10 min and washed three times with PBS + 75 mg/ml Glycine (pH=7.4, Sigma). Unspecific binding sites were blocked for 1 hour at RT with 10% goat serum in IF-wash solution (PBS + 5 mg/ml NaN^3^, 10 mg/ml bovine serum albumin, 2% TritonX-100 and 0.4% Tween20, pH=7.4, Sigma). Afterwards, primary antibodies in blocking solution were added and incubated at 4°C overnight. The next day, cells were washed 3x with IF-wash and then incubated with fluorescently labeled secondary antibodies in blocking solution for 1 hour at RT if primary antibodies were unlabeled. Subsequently, cells were washed 3x with IF-wash and 2x with PBS and then incubated in PBS + 1 µg/ml Hoechst for 20 min at RT. Cells were again washed with PBS, removed from the surface and transferred into 8-well Nunc™ Lab-Tek™ Chamber Slides (ThermoFisher) for improved fluorescence detection. The following antibodies were used in this study: Rabbit anti-Vimentin antibody Alexa Fluor^®^ 594 (1:100, EPR3776, abcam), mouse anti-β-Actin antibody (1:200, 8H10D10, Cell Signaling), Mouse anti-Cytokeratin 15 antibody (1:50, LHK15, ThermoFisher), Goat anti-mouse Alexa Fluor^®^ 594 (1:200, Cell Signaling).

3×3 images per well (20 Z-stacks per position) were acquired automatically on a Zeiss LSM780 Axio Observer confocal microscope equipped with a 10x/0.3 air objective (Zeiss EC PLAN-NEOFLUAR) using a custom Zeiss VBA macro. Beside brightfield images, lasers and filters were set to measure fluorescence emitted from Hoechst (DNA) and from Alexa Fluor^®^ 594-labeled antibodies. Images were analyzed using a custom KNIME workflow in which protein abundances per classified spheroid were defined as mean pixel intensity of the fluorescence signal emitted from labeled antibodies.

#### RNA FISH

For histological preparation, MCF10CA spheroids were cultured and isolated as described above, fixed in 2% Formaldehyde solution for 20 min at RT and washed twice with PBS. Afterwards, spheroids were incubated in PBS + 15% sucrose (Sigma) and PBS + 30% sucrose (both 15 min at RT), embedded in Richard-Allan Scientific™ Neg-50™ Frozen Section Medium (ThermoFisher) and frozen in the gaseous phase of liquid nitrogen.

For the same purpose, ‘big’ (70-100 µm) and ‘small’ (70-100 µm) colon tumor spheroids derived from single-cells were isolated with (reverse-) filtering as described above. This step was added for histological preparation in order to distinguish between small spheroids and big spheroids that were sliced in peripheral regions. Spheroids were then fixed with 4% Formaldehyde solution for 20 min at 4°C, washed twice with PBS and incubated in 30% sucrose at 4°C overnight. The next day, spheroids were embedded in Neg-50™ and frozen in the gaseous phase of liquid nitrogen.

For both cultures, sectioning was performed at −20°C on a cryostat (Leica) and 10 µm slices were mounted on Superfrost Plus slides (ThermoFisher). Embedded specimens and cryosections were stored at −80°C until further use. For highly sensitive RNA fluorescence in-situ hybridization (RNA-FISH), we employed the RNAscope^®^ Fluorescent Multiplex Assay 2.0 (ACDbio). Cryosections were processed as described in the ‘Sample Preparation Technical Note for Fixed Frozen Tissue’ and the ‘Fluorescent Multiplex Kit User Manual PART 2’. Briefly, cryosections were pretreated with Protease IV (ACDbio) for 15 min at RT. Afterwards, transcript-specific probes were hybridized at 40°C for 90 min followed by stepwise hybridization of probes for signal amplification and fluorescent detection (Amp-1-FL – Amp-4-FL). Up to three transcripts were labeled by Alexa488, Atto550 and Atto647 fluorescent dyes. Following mRNA targeting probes were used: SNAI2 (Alexa488, #554581), DEFA5 (Alexa488, #423981), MYC (Atto550, #311761-C2), CD44 (Atto647, #311271-C3), TFF3 (Alexa488, #403101), PROX1 (Atto550, #530241-C2). Finally, cryosections were counterstained with DAPI, mounted in SlowFade™ Gold Antifade solution (ThermoFisher) and stored at 4°C until further use.

RNA-FISH images were acquired on a Leica SP8 confocal laser-scanning microscope equipped with a 40x/1.30 oil objective (Leica HC APO CS2). Images of individual spheroids at 1024×1024 pixel resolution were generated semi-automatically using the ‘Mark and Find’ option in the Leica SP8 acquisition software. To cover the whole 10 µm cryosection height, a Z-range of 20 µm was acquired by 15 stacks (1.43 µm distance between frames). Lasers and filters were set to match fluorescent properties of DAPI and abovementioned dyes. MCF10CA spheroids were already classified as ‘aberrant’ or ‘round’ manually during imaging, whereas colon tumor spheroid classes were already separated during sample preparation. For analysis of RNA-FISH imaging data we used a custom KNIME workflow in which we defined the relative transcript expression per spheroid as quantified pixel percentage that exceeds a calculated background threshold per spheroid.

#### Cell count determination by light sheet imaging and 3D segmentation

To estimate the cell number in spheroids in HT-pheno-seq experiments based on acquired images, we generated a high-resolution 3D reference dataset to determine the linear relationship of size (area) and cell count. CRC spheroids were stained with 1 µg/ml Hoechst and CellTracker Red CMPTX (10 µM) for 3 hours and isolated and fixed as described above. Subsequently, spheroids were mounted in 2% low-melting agarose (Sigma) and 3D images were acquired using a Dual-View Inverted Selective Plane Illumination Microscope (ASI di-SPIM) using Nikon 40x/0.80W NA NIR-Apo water dipping objectives. Dual view raw data was processed to generate isotropic images at a resolution of 0.325px/µm (400 images per Z-stack, 0.325 µm distance). Pre-processing and 3D segmentation were performed with a custom KNIME workflow. Cell counts and corresponding 2D spheroid dimensions were exported from KNIME and further analyzed and plotted using a custom R script. Briefly, we first converted the respective pixel numbers for minor and major axis of all 2D spheroid masks to metric distances. Facing the challenge of variable spheroid morphologies, we further approximated the spheroid size as the product of its biggest and smallest diameter (i.e. the minor and major axis). Subsequently, we plotted the approximated size of every spheroid against its respective cell count determined by 3D segmentation in KNIME. A linear model was fitted through the points and the obtained slope was used to calculate cell number estimations for HT-pheno-seq experiments.

### Sequencing data analysis

#### Pre-processing of RNA-seq data and library quality control

An automated in-house workflow was established for single-cell and spheroid RNA-seq data pre-processing. Briefly, short read quality was evaluated using FastQC. For iCELL8 libraries, barcodes from the first 21 bp read were assigned to the well of origin with the Je demultiplexing suite. Cutadapt was used to trim remaining primer sequences, Poly-A/T tails and low-quality ends (<25). In addition, since NextSeq (Illumina) encodes undetected base as incorrect ‘G’ with high quality, Cutadapt’s ‘—nextseq-trim’ option was used for correct quality trimming. Trimmed reads were mapped to the reference genome hs37d5 (1000 genomes project) using STAR aligner. Mapped BAM files were quantified using featureCounts with gencode v19 as reference annotation.

RNA-seq libraries that did not match the following criteria were filtered out: MCF10CA scRNA-seq: (i) > 300,000 reads, (ii) > 3000 detected genes (i.e. >0 read count), (iii) < 10% mitochondrial reads; MCF10CA pheno-seq: (i) > 100,000 reads, (ii) > 2000 detected genes, (iii) < 15% mitochondrial reads; Colon spheroid pheno-seq:(i) > 200,000 reads, (ii) > 3000 detected genes, (iii) < 15% mitochondrial reads. In order to compare the performance of scRNA-seq and pheno-seq methods in detecting genes, MCF10CA sequencing libraries were downsampled to 100,000 reads by a custom R script.

Wells/Spheroids with imaging artifacts (e.g. segmentation errors) were removed if detected during combined downstream analysis.

#### RNA-seq subpopulation and differential expression analysis

To identify expression signatures that separate distinct cellular subpopulations, we analyzed transcriptional heterogeneity by pathway and gene set overdispersion analysis (PAGODA/SCDE-package^24^). First, genes with less than 10 mapped reads in the whole dataset were not considered for further analysis. Next, PAGODA constructs error models for individual cells using a binominal/Poisson mixture model, thereby controlling for technical aspects of variability, like effective sequencing depth, drop-out rate and amplification noise. For K-nearest neighbor error modelling, k was set to 30 (except for the manual pheno-Seq dataset: k=3), and the minimum number of reads required to be considered non-failed was set to 2. Afterwards, PAGODA performs weighted principal component analysis (wPCA) on annotated and *de-novo* identified gene sets in order to identify those that exhibit statistically significant variability. Generally, the scores for the first principal component are presented if not stated otherwise. Annotated hallmark (H) and gene ontology (GO_C5) gene sets were derived from the Molecular Signature Database (MSigDB). *De-novo* gene sets were identified by hierarchical clustering (Ward method; dendrogram was cut into 150 clusters). Pathway overdispersion was calculated as Z-score relative to the genome-wide model and corrected Z-scores (cZ) were computed using multiple hypothesis testing using the Holm procedure. Hierarchical clustering is then performed on the top significant aspects of heterogeneity and redundant aspects of heterogeneity were grouped with a similarity threshold of 0.7. Up to 10 top significant aspects were used for visualization. In addition, 2D t-SNE maps^54^ were generated based on PAGODA’s weighted Pearson correlation distances. Finally, the following confounding expression signatures (e.g. technical aspect or cell cycle influence) were removed using the ‘pagoda.substract.aspect’ function:

1. For all datasets we corrected for the influence of gene coverage (estimated as a number of genes with non-zero magnitude per cell)
2. MCF10CA scRNA-seq: GO_REGULATION_OF_CELL_CYCLE and HALLMARK_G2M_CHECKPOINT;
3. MCF10CA HT-pheno-seq: GO_ NUCLEOSIDE_MONOPHOSPHATE_ METABOLIC _PROCESS, GO_MITOCHONDRIAL_ENVELOPE, GO_STRUCTURAL _MOLECULE _ACTIVITY, GO_ HOMEOSTATIC_PROCESS and associated *de-novo* identified gene sets.

Differentially expressed genes (MCF10CA: fold change > 1.3; adjusted p-value < 0.1; CRC: fold change > 1.5; adjusted p-value < 0.05) between detected subpopulations that refer to observed visual phenotypes (k-means clustering, k=2) were identified by the SCDE-package^26^.

### In-silico reconstruction of synthetic pheno-seq expression profiles from single-cell data

Synthetic spheroid expression profiles were reconstructed from scRNA-seq data by randomly dividing cells either derived from round and aberrant phenotypes in four groups each in four independent randomizations. Read counts for each gene were then averaged over each group, resulting in eight synthetic spheroid profiles (4 round and 4 aberrant) that were then analyzed by PAGODA similar to the manual pheno-seq dataset.

### Deconvolution of the CRC spheroid dataset by maximum likelihood inference

In order to infer heterogeneous regulatory states informative for single cell expression by deconvolution, we adapted a maximum likelihood inference approach initially developed to identify cell-to-cell heterogeneities from random 10-cell samples^41^ (Stochastic Profiling, Fig. 3a). Here, we allowed each sample to consist of different numbers of cells (implemented in the R package stochprofML version 2.0: https://github.com/fuchslab/stochprofML)

The algorithm assumes that the expression of a spheroid linearly scales with its cell number. We approximated absolute counts per spheroid by using estimated cell numbers derived from light sheet microscopy and image analysis: First, counts per spheroid were divided by the respective estimated cell number, and the minimal average mRNA count per cell was determined (2374.644). Afterwards we downsampled the whole dataset to 2300 counts per cell resulting in a perfect correlation of mRNA counts and cell numbers. (Supplementary Figure 9b). The downsampled dataset was filtered by removing genes with less than one count per well on average over the original CRC spheroid dataset and genes with less than 5 counts in at least two wells, leaving 13,868 genes that are taken into account during the profiling procedure. To avoid problems with zeros and log-normal distributions, all zeros were transformed to 0.1.

### Statistical analysis and visualization

Statistical analysis and visualization of sequencing data was done in R (Version 3.3.1) or R studio (https://www.rstudio.com/) using PAGODA/SCDE, ggplot2, ComplexHea™aps^55^, the stats package (R version 3.3.1), stochprofML (R version 3.4.1) and in Graph Pad Prism 7 (https://www.graphpad.com/scientific-software/prism/). Gene set enrichment analysis was done by computing overlaps between identified class-specific signatures and gene sets derived from the Molecular Signature Database^25^ (MSigDB, https://software.broadinstitute.org/gsea/msigdb).

## Data and code availability

Raw sequencing data for MCF10CA are accessible at the European Nucleotide Archive (https://www.ebi.ac.uk/ena) under Accession Number PRJEB26737.CRC HT-pheno-seq raw sequencing data have been deposited at the European Genome-Phenome Archive (http://www.ebi.ac.uk/ega/) under Accession Number EGAS00001002999.

All KNIME image analysis workflows, R code for PhenoSelect and PAGODA/SCDE RNA-seq analysis as well as a download link for MCF10CA HT-pheno-seq image data with all necessary components to run the pre-processing workflow and/or the PhenoSelect web app can be found in the pheno-seq github repository (https://github.com/eilslabs/pheno-seq). Information on the automated in-house RNA-seq workflow is available upon request. The newest version of stochProfML 3.4.1 can be found under: https://github.com/fuchslab/stochprofML.

## Supplementary files and data

Supplementary text and figures as well as a step-by-step HT-pheno-seq protocol are provided as single supplementary PDF file.

